# Intermodal consistency of whole-brain connectivity and signal propagation delays

**DOI:** 10.1101/2023.08.28.555074

**Authors:** Maciej Jedynak, Emahnuel Troisi Lopez, Antonella Romano, Viktor Jirsa, Olivier David, Pierpaolo Sorrentino

## Abstract

Measuring the propagation of perturbations across the human brain and their transmission delays is critical for network neuroscience, but it is a challenging problem that still requires cross-validation approaches. Here, we compare results from a recently introduced, non-invasive technique of functional delays estimation from source-reconstructed electro/magnetoencephalography, to the corresponding findings from a large dataset of cortico-cortical evoked potentials estimated from intracerebral stimulations of epileptic pharmaco-resistant patients. The two methods yield significantly similar probabilistic connectivity maps and signal propagation delays. This similarity suggests a correspondence between the mechanisms underpinning the propagation of spontaneously generated scale-free perturbations (i.e. neuronal avalanches observed in resting state activity studied using magnetoencephalography) and the spreading of cortico-cortical evoked potentials. This manuscript provides evidence for the accuracy of a subject-specific estimate of functional delays obtained non-invasively from reconstructed sources. Conversely, our findings show that estimates obtained from externally-induced perturbations capture physiological activities. In conclusion, this manuscript constitutes a cross-validation between two different modalities. Importantly, the capability to measure delays non-invasively (as per MEG) paves the way for the inclusion of functional delays in personalized large-scale brain models as well as in diagnostic and prognostic algorithms.

## Introduction

Brain functions are thought to be emergent from the interaction of multiple brain areas, which is fine-tuned and generates complex dynamics^1^. This complexity is informative both in physiological and pathological conditions at rest and in the context of brain perturbations, such as pulses or pharmacological manipulations. For example, quantitative tools based on external perturbations of the brain have shown that the complexity of brain dynamics relaxation after stimulation predicts the level of consciousness ^2^. Even at rest, large-scale activities self-organize in dynamical, aperiodic, large-scale nonlinear bursts, and, in disease, such burst dynamics are simplified and stereotyped ^2,3^. Hence, the elements that underpin ‘healthy” dynamics are essential, both from a theoretical and experimental perspective ^4^. Recent evidence showed that the scaffolding of the white-matter bundles linking gray matter regions partly determines the spatial spread of burst dynamics observed *in vivo*^5^. In fact, the probability of two brain regions activating sequentially is proportional to the coupling intensity along the brain tract linking them ^5^. Subsequently, personalized large-scale brain models typically couple the equations according to the structural connectome ^6^. However, theoretical arguments highlight that the influence of the connectome is not limited to the spatial evolution of large-scale dynamics, but also determines its temporal evolution, by affecting the functional delays across brain regions ^7^. In large-scale modeling, it is often assumed that the functional delays are proportional to the tract length alone ^4^. This simplifying assumption is typically made given the difficulty of measuring the functional delays *in vivo* across the whole brain. Nevertheless, two different approaches have been introduced recently in order to measure functional delays in vivo.

The first approach, based on electro/magnetoencephalography (E/MEG), relates to the presence of correlates of critical dynamics (i.e. neuronal avalanches) during resting-state activity ^8^. Brain activities were estimated by source-reconstructing resting-state, eyes closed MEG data, filtered between 0.5 and 48 Hz, acquired from 58 healthy participants (see methods). Neuronal avalanches are aperiodic, scale-free bursts of activations that characterize the connectivity during resting state. The way such bursts spread in space and time can be conveyed using the recently-described avalanche transition matrix (ATM). For tracts that are known to have direct structural connections, it is possible to estimate the time it takes a neuronal avalanche to recruit any two consecutive regions ^9^. In this way, one can estimate functional delays. This approach has been shown to successfully capture longer delays along white matter tracts that were affected by demyelinating lesions in multiple sclerosis patients ^9^.

The second approach is based on a functional tractography project (F-TRACT, https://f-tract.eu) which is based on a group analysis of a large cohort of patients suffering from pharmaco-resistant focal epilepsies who were implanted with intracerebral stereoencephalographic (SEEG) electrodes^10,11^. Those electrodes were used to deliver single pulse electrical stimulations and to record resulting responses, cortico-cortical evoked potentials (CCEP). By merging the data from the whole cohort of patients, one can compute the probability that stimulation to a certain brain region will evoke a CCEP in any other region ^12,13^. Moreover, the time latency (delay) needed for propagation between the two regions can be directly measured. Here, by aggregating stimulations performed in 583 patients, it was possible to retrieve a whole-brain atlas of such probabilities and delays ^14,15^ (see Materials and Methods).

In this manuscript, we aimed to compare probability and delay measures obtained with the two above-mentioned methods, providing cross-validation for both. We assume that a high degree of similarity between the results obtained from the two methods would support the hypothesis that both the spreading of the SEEG-evoked perturbations and of the neuronal avalanches are two facets of the same, universal large-scale communication mechanisms. A high degree of similarity would mean that the probabilistic map obtained from the perturbative SEEG method could be considered a good estimate for connectivity spontaneously occurring during resting state (as observed using E/MEG). Conversely, the probability and delay measurements obtained from the non-invasive E/MEG method, based on the spread of avalanches, could be considered accurate estimates, since they are in accordance with the measurements observed when delivering stimulations directly via implanted electrodes. As a consequence, a new tool for noninvasive derivation of brain connectivity and communication delays could be incorporated into relevant diagnostics procedures in E/MEG. The results presented in this paper are based on MEG data.

## Results

We present a comparison of two connectivity maps of the human brain, each independently obtained from two different modalities and analysis methods applied to two different cohorts of subjects. The first map contains probabilistic functional connections between any two brain parcels as defined by the AAL parcellation ^16^. Connections are defined as the probability of observing the propagation of activity from the source to the target parcel. The second map informs about the time elapsed during such propagation. The two compared modalities are: 1) a non-invasive method based on the merging neuronal avalanches from source-reconstructed MEG and structural data from 44 healthy subjects and 2) functional tractography performed on a large dataset of nearly six hundred patients who underwent intracerebral implantations with SEEG electrodes utilized to deliver electrical stimulations and record the evoked brain-wide responses.

We first present the comparison of probabilities (for observing a significant connection between a pair or parcels). They are shown in Figure 1 for the MEG dataset (top row, left) and for the F-TRACT dataset (top row, middle). In both cases, the source parcel is found along the ordinate axis and the target parcel along the abscissa. A scatter plot relating those two matrices is shown in Figure 2 (top, right), demonstrating that there is a positive linear correlation between the results from the two datasets. This is confirmed by a high value of Pearson correlation r=0.58, p<0.001.

**Figure 1.**
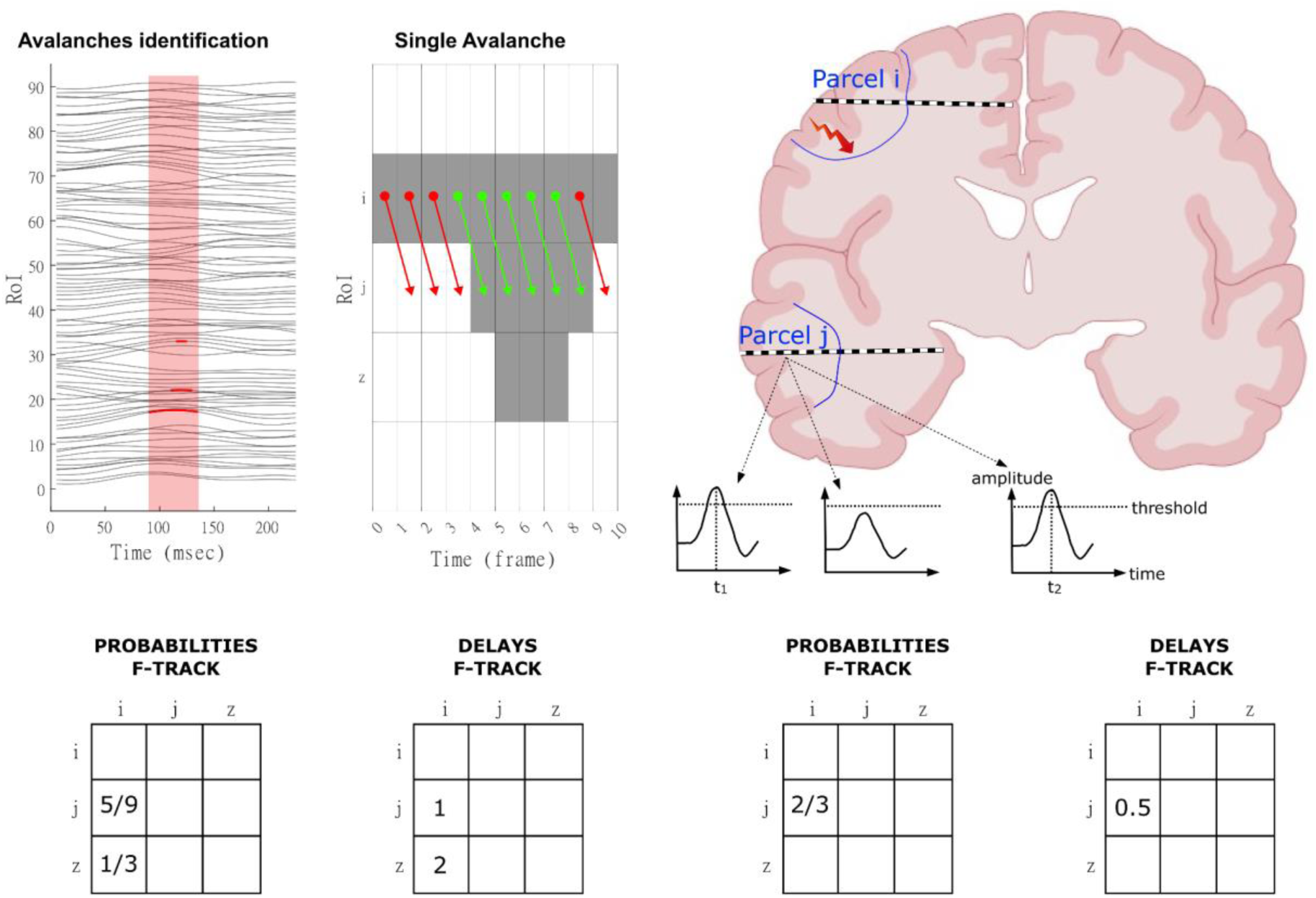
**Left.** Source-reconstructed MEG data are reported at the top-left. The data is z-scored and thresholded. The probabilities are defined as the frequency with region *j* went above the threshold after region *i* did. Finally, the delays are defined as the median time it took region *j* to go above the threshold after region *i* did. **Right.** Schematic representation of the CCEP processing. In the toy example reported, three stimulations have been delivered to parcel *i* and recorded from parcel *j*, via intracranial electrodes depicted here with dashed thick lines. The time courses represent z-scored (with respect to pre-stimulus baseline) responses to stimulation. The transition probabilities for the tract *i,j* are defined as the percentage of pulses delivered that elicited an above-threshold response in the recording electrode. In the toy example depicted in the figure, two out of the three stimulations elicited an above-threshold response. Hence, the corresponding probability equals ⅔. With respect to the delays, we have computed the median time it took from the electrical pulse delivery to the maximum of the first CCEP peak appearing after the threshold-crossing in the receiving electrode. In the toy example, the delays for the two pulses that reached the threshold correspond to 1 and 2, resulting in a median of 0.5.

**Figure 2.**
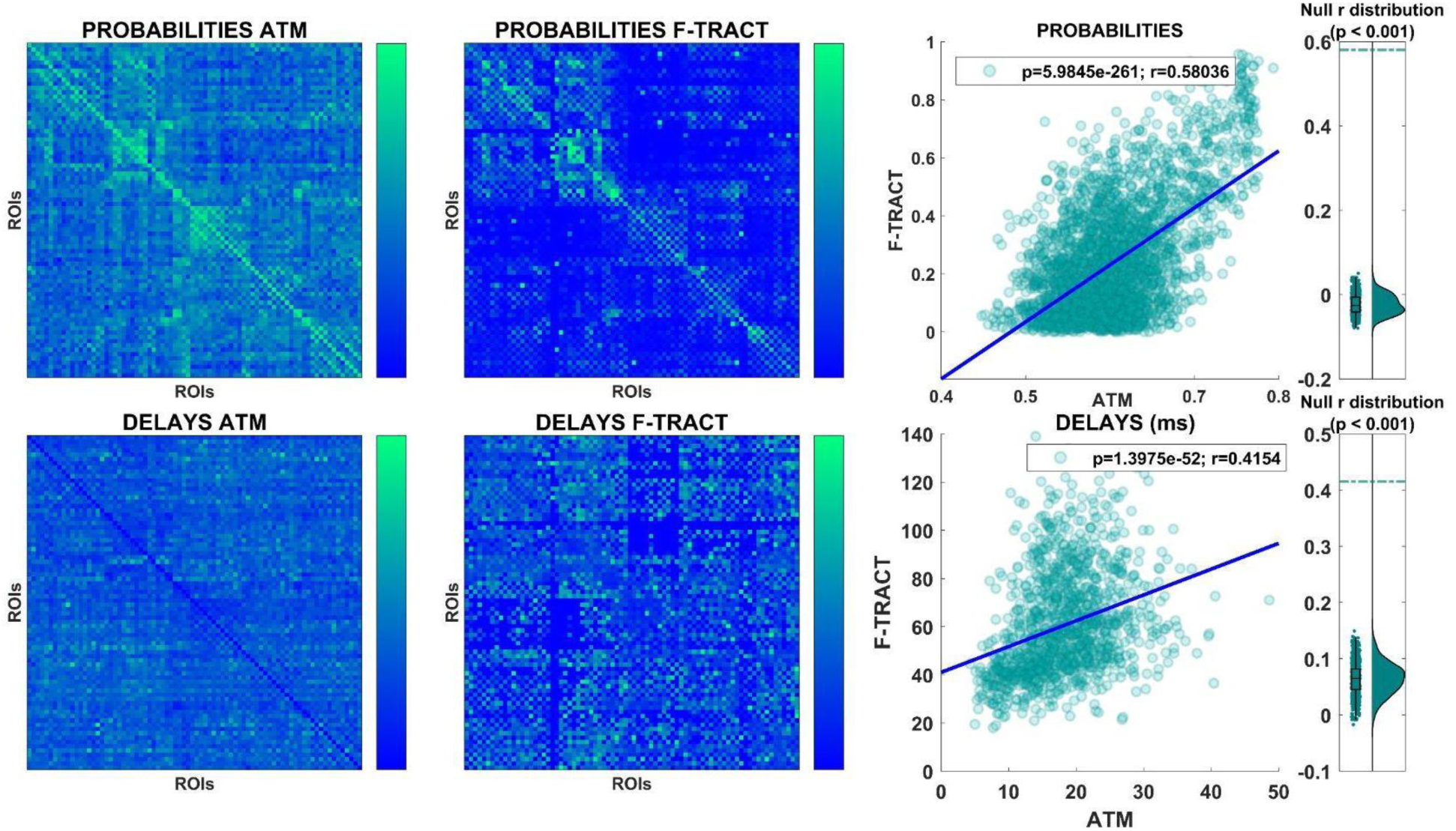
**Top-left:** Connectivity probabilities as obtained from the Avalanche Transition Matrices (ATM). Source ROIs are in rows, and target ROIs are in columns. This convention remains valid for the remaining presented connectivity matrices. **Top-center:** The matrix contains transition probabilities as obtained from the F-TRACT dataset. **Top-right:** each dot of the scatterplot corresponds to a connection between two parcels, the x-axis informs about probabilities obtained using the ATMs, and the y-axis about probabilities as obtained using F-TRACT. The least-squares fit line is also reported. To the far right, the distribution of the correlations was obtained by shuffling the temporal course of the ATMs (while preserving their spatial properties - see methods for the details). The green line corresponds to the empirically-observed correlation. **Bottom-left:** the matrix contains the delays as obtained from the F-TRACT dataset. **Bottom-center:** the matrix contains the delays as obtained from the ATMs. **Bottom-right:** similarly as above, each dot of the scatterplot corresponds to a connection between two ROIs, the x-axis shows the delays obtained using the ATMs, and the y-axis the delays obtained using F-TRACT. The least-squares line is also reported. To the far right, the distribution corresponds to the correlations obtained with the surrogate data, and the green dashed line marks the empirically-observed correlation.

To test for the likelihood of obtaining the observed correlation by chance, we proceeded with surrogate analysis as follows. Similarly to ^5^, we randomly shuffled the temporal sequence of activations within each MEG avalanche, while keeping the spatial structure unaffected. New avalanche transition matrices were estimated based on the shuffled avalanches and then related to the original F-TRACT-based probabilities. This procedure was repeated one thousand times, yielding a null distribution to be expected given random dynamics with preserved spatial structure. Our results presented in Figure 2, distributions to the right confirm that obtaining the observed correlation given random dynamics is highly unlikely. Then, we moved on to compare delays in the two datasets. Again, as shown in Figure 2, bottom row, the delays estimated from the two datasets also appear to be correlated (p<<0.001, r=0.42), confirming that both methods capture comparable temporal dynamics in terms of the propagation of the perturbations. Similarly, here, the surrogate analysis confirmed the validity of our results (p< 0.001).

## Discussion

In this manuscript, we set out to compare the properties of large-scale functional connectivity as measured in two different modalities. Specifically, the compared maps were: probabilistic connectivity and delays related to stimulus propagation from the source to the target brain parcel. Each map was independently computed in two considered modalities.

First, probabilities and delays were provided by a recently developed technique extracting neuronal avalanches from MEG recordings. Then, the probabilities for connections and the corresponding delays were also estimated from F-TRACT, a large dataset of intracerebral electric brain stimulations of the human brain. Our results show that both the probabilities and the functional delays are correlated between the modalities, thereby suggesting the validity of both techniques. However, the two modalities are very different from a technical standpoint, and they are derived from different cohorts. The probabilities in the two considered datasets express different measures, therefore we do not expect agreement of absolute values but rather a correlation between them. In particular, in order to compute the transition probabilities in the ATMs, only the avalanches that spread across a given edge are taken into account to estimate the transition probability of that edge. On the other hand, in the F-TRACT the responses of a given region are compared to the total of the performed stimulations. This might explain the fact that the transition probabilities in the ATM remain higher (above ∼0.45) as compared to the ones estimated from the F-TRACT. Similarly, the maximal delay measured in the ATM technique is 64 ms, whereas longer delays were observed in F-TRACT (presumably due to polysynaptic connections). Moreover, the F-TRACT dataset could be considered the gold standard, given the fact that the electrodes are implanted directly within the brain, which provides better spatial resolution compared to source-reconstructed MEG. Furthermore, with respect to the delay estimation, the F-TRACT dataset offers a well-controlled setting because the SEEG stimulation procedure allows one to know the exact moment and characteristic of the stimulation, as well as to register a defined pulse at the recording electrodes^15^. Nevertheless, the drawback of this approach lies in the fact that direct electrical stimulation differs from physiological brain activity. In fact, the amplitude of the stimulus delivered, typically in the order of 1-5 mA, is far more intense than typical neuronal inputs. Moreover, although electrodes with high rates of interictal epileptic-like spiking were excluded from the analyses, we cannot rule out that the remaining recordings in the F-TRACT dataset might be partly affected by the pathology.

This is different from the MEG dataset, which gathers data from healthy subjects. The source-reconstructed MEG dataset estimates spontaneously propagating perturbations with a different experimental setup and data analysis pipeline, making it unlikely that similar results might be spurious or due to a particular technique. In fact, source reconstruction in E/MEG is an ill-posed problem ^12^: the estimated source activity holds a degree of uncertainty, which drastically rises for the estimation of deep sources. Furthermore, applying the framework of criticality, and, in particular, that of neuronal avalanches, allows us to track spontaneous perturbations that occur in the brain on the large-scale ^5,8^. In this sense, one tracks scale-free perturbations spreading spontaneously on the large scale, avoiding potential confounds induced by external stimulations. Although obtained from two independent cohorts, results from the two modalities are highly similar, suggesting that the F-TRACT dataset could be interpreted in the context of resting state in healthy individuals, because of the two following reasons. First, there is a correlation with healthy individuals despite the bias related to epilepsy. Second, F-TRACT data are obtained in a stimulation paradigm, yet they resemble data obtained during resting state. Conversely, our result suggests the reliability of the non-invasive estimation of delays and probabilities obtained by tracking neuronal avalanches from resting-state, source-reconstructed M/EEG data.

In conclusion, our manuscript cross-validates two different techniques for the estimation of connection probabilities and conduction delays from whole-brain neurophysiological data. The relevance of these results is two-fold. On the one hand, delays estimated non-invasively might extend the personalization of large-scale models such as The Virtual Brain ^9^. In fact, the state-of-the-art models assume delays as scaling proportionally to white matter tract lengths. The subsequently estimated delays range differently as compared to the observed ones ^9^ Similarly, these models assume all connections to be symmetric, as directionality is not known from diffusion MRI. Nevertheless, both modalities discussed in this paper provide information about connection directionality. The personalized modeling approach could be improved with the here-considered noninvasive method of delay and directionality estimation at the single-subject level. Furthermore, estimating the damage of the functional delays holds promise to improve the diagnostic and prognostic management of patients with neurological and, potentially, psychiatric ailments.

## Methods

### MEG dataset

#### Participants

Fifty-eight right-handed and native Italian speakers were considered for the analysis. To be included in this study, all participants had to satisfy the following criteria: a) to have no significant medical illnesses and not to abuse substances or use medications that could interfere with MEG/EEG signals; b) to show no other major systemic, psychiatric, or neurological illnesses; and c) to have no evidence of focal or diffuse brain damage at routine MRI. The study protocol was approved by the local Ethics Committee. All participants gave written informed consent.

#### MRI acquisition

Diffusion MRI data were acquired for the same individuals using a 1.5 Tesla machine (Signa, GE Healthcare). Preprocessing was performed using the software modules provided in the FMRIB Software Library (FSL, http://fsl.fmrib.ox.ac.uk/fsl). Data were corrected for head movements and eddy current distortions using the “eddy_correct” routine, rotating diffusion sensitizing gradient directions accordingly, and a brain mask was obtained from the B0 images using the Brain Extraction Tool routine. A diffusion-tensor model was fitted at each voxel, and fiber tracks were generated over the whole brain using deterministic tractography, as implemented in Diffusion Toolkit (FACT propagation algorithm, angle threshold 45°, spline-filtered, masking by the FA maps thresholded at 0.2). For tractography analysis, the ROIs of the AAL atlas and of a MNI space-defined volumetric version of the Desikan-Killiany-Tourville (DKT) ROI atlas were used, both masked by the GM tissue probability map available in SPM (thresholded at 0.2). To this end, for each subject, FA volumes were normalized to the MNI space using the FA template provided by FSL, using the spatial normalization routine available in SPM12, and the resulting normalization matrices were inverted and applied to the ROIs, to apply them onto each individual. The quality of the normalization was assessed visually. For each individual, the number of streamlines interconnecting each pair of regions was enumerated using custom software written in Interactive Data Language (IDL, Harris Geospatial Solutions, Inc., Broomfield, CO, USA). Results were replicated using both the AAL and the DKT atlases. In supplementary analyses, connectomes were also mapped using diffusion MRI data for 200 participants from the Human Connectome Project using an alternative workflow. The resulting individual connectomes were then averaged to yield a group-consensus connectome. Further details are available in SI. See Fig.1 for an overview of the methods.

#### MEG acquisition and pre-processing

MEG pre-processing and source reconstruction were performed as in^18^. In short, the MEG registration was divided into two eyes-closed segments of 3:30 minutes each. To identify the position of the head, four anatomical points, and four position coils were digitized. Electrocardiogram (ECG) and electrooculogram (EOG) signals were also recorded ^19^. The MEG signals, after an anti-aliasing filter, were acquired at 1024 Hz, then a fourth-order Butterworth IIR band-pass filter in the 0.5-48 Hz band was applied. To remove environmental noise, measured by the reference magnetometers, we used Principal Component Analysis ^20^. We adopted Independent Component Analysis to clean the data from physiological artifacts ^21^, such as eye blinking (if present) and heart activity (generally one component). Noisy channels were identified and removed manually by an expert rater. 47 subjects were selected for further analysis. The time series of neuronal activity were reconstructed based on the Automated Anatomical Labeling (AAL) ^22,23^. To do this, we used the Linearly Constrained Minimum Variance (LCMV) beamformer algorithm based on the native MRIs^17^ . Finally, we excluded the ROIs corresponding to the cerebellum because of their low reliability in MEG. However, when these regions were included, the results were replicated. All the preprocessing and the source reconstruction were performed using the Fieldtrip toolbox ^24^.

#### Transition matrices

Each source reconstructed signal was binned (such as to obtain a branching ratio ∼1, see SI) and then z-scored and binarized, such that, at any time bin, a z-score exceeding ± 3 was set to 1 (active); all other time bins were set to 0 (inactive). An avalanche was defined as starting when any region is above threshold, and finishing when no region is active, as in ^3^. The results reported refer to a binning =3, corresponding to a branching ratio of 1. An avalanche-specific transition matrix (TM) was calculated, where the element (*i*, *j*) represented the probability that region *j* was active at time *t+ẟ*, given that region *i* was active at time *t*, where *ẟ*∼3ms. The TMs were averaged per participant, then per group, and finally symmetrized^5^.

#### Construction of random surrogates

Randomized transition matrices were generated to ensure that associations between transition probabilities and structural connectivity could not be attributed to chance. Avalanches were randomized across time, without changing the order of avalanches at each time step. We generated a total of 1000 randomized transition matrices and the Spearman rank correlation coefficient was computed between each randomized matrix and the matrices derived from the F-TRACT dataset. This yielded a distribution of correlation coefficients under randomization. The proportion of correlation coefficients that were greater than, or equal to, the observed correlation coefficient provided a p-value for the null hypothesis that the structure of the avalanches is not related to the spreading of the cortico-cortical evoked potential as recorded in the F-TRACT.

#### Estimation of delay matrices from neuronal avalanches

The delays were estimated for each avalanche, as in ^9^. In an avalanche, from the moment region *i* activated, we recorded how long it took region *j* to activate. These are what we considered to be delays. Hence, for each avalanche, we obtained a matrix, in which the rows and columns represented brain regions and the entries contained the delays. We then averaged across all the avalanches belonging to one subject, obtaining an average *ij^th^* delay. The average was performed disregarding zero entries since each avalanche-specific matrix is very sparse. With this procedure, a subject-specific delay matrix was built. Averaging across subjects (again discarding zero entries) yielded a group-specific matrix.

### F-TRACT dataset

SEEG recordings analyzed in this study come from the F-TRACT project (https://f-tract.eu) ^13–15^ that to this day gathered data from over one thousand pharmaco-resistant epileptic patients, who in the course of preparation for the brain resection surgery underwent intracerebral implantation^10,11^. Since the MEG cohort consists of only adults, we selected from the F-TRACT dataset only patients being at least 18 years old. As a result, in the analysis, we considered 584 implantations (288 male, 291 female, 5 unspecified) from 573 unique patients (283 male, 285 female, 5 unspecified) coming from 21 centers. The average age was 33 with sd 10. In total 2,786,513 recordings were considered, following 29,869 stimulations (73% biphasic, 27% monophasic) with 85 electrodes per stimulation on average, mean stimulating current intensity of 3.3 mA, and pulse width 1 ms.

Derivation of the connectivity atlas is described in detail in ^14^. Briefly, the procedure was as follows. In order to minimize potential epileptic effects, for each subject we discarded 20% of recordings having the highest interictal spike count rate as assessed with DELPHOS software ^25,26^. To mitigate volume conduction effects, we re-referenced recorded signals to bipolar montage. The signals were band-pass filtered in the range 1 and 45 Hz and responses to all pulses in a stimulation were averaged. In order to account for individual and local specificity, we z-scored the signal using the mean and standard deviation of spontaneous fluctuations computed on the pre-stimulation interval [-400, -10 ms]. Responses to stimulation trespassing z-score threshold Z=5 within a 200 ms post-stimulation time window were considered significant. Delays were obtained from significant responses as the latency passed from stimulation until the occurrence of the first peak maximum after the threshold crossing. For both probabilities and delays, we only consider values computed from at least 100 measurements.

AAL parcellation ^17^ allocation to SEEG electrode contacts was performed from their MNI coordinates estimated after the MNI normalization of the preoperative MRI of individual subjects, which were first co-registerred with post-operative MRI or CT-scans showing implanted electrodes. On the group level, we aggregated all recordings having the same source and target parcels and we derived frequentist probability for a connection as the ratio of the number of responses considered significant over the number of stimulations between these two parcels. Similarly, the group-level delay was estimated by the median value from delays characterizing significant responses.

## Notes

### Competing Interest Statement

The authors have declared no competing interest.

